# Building mathematical models of biological systems with modelbase, a Python package for semi-automatic ODE assembly and construction of isotope-specific models

**DOI:** 10.1101/362954

**Authors:** O. Ebenhöh, M. van Aalst, N.P. Saadat, T. Nies, A. Matuszyńska

## Abstract

The `modelbase` package is a free expandable Python package for building and analysing dynamic mathematical models of biological systems. Originally it was designed for the simulation of metabolic systems, but it can be used for virtually any deterministic chemical processes. `modelbase` provides easy construction methods to define reactions and their rates. Based on the rates and stoichiometries, the system of differential equations is assembled automatically. `modelbase` minimises the constraints imposed on the user, allowing for easy and dynamic access to all variables, including derived ones, in a convenient manner. A simple incorporation of algebraic equations is, for example, convenient to study systems with rapid equilibrium or quasi steady-state approximations. Moreover, `modelbase` provides construction methods that automatically build all isotope-specific versions of a particular reaction, making it a convenient tool to analyse non-steady state isotope-labelling experiments.

## (1) Overview

### Introduction

Well designed mathematical models are excellent theoretical frameworks to analyse and understand the dynamics of a biological system. Here, the design process itself is the first important scientific exercise, in which biological knowledge is collected, organised and represented in a new, systematic way, that allows defining the model assumptions and formulating them in the language of mathematics. A working model then enables testing new hypotheses and allows for novel predictions of the system’s behaviour. Kinetic models allow simulating the dynamics of the complex biochemistry of cells. Therefore, they have the power to explain which processes are responsible for observed emergent properties and they facilitate predictions on how the system behaves under various scenarios, such as changed environmental conditions or modification of molecular components. Unfortunately, the construction of mathematical models is often already a challenging task, hampered by the limited availability of measured physiological and kinetic parameters, or even incomplete information regarding the network structure. It is therefore highly desirable to make the overall process of model construction as easy, transparent and reproducible as possible. Providing a toolbox with a wide range of methods that flexibly adapt to the scientific needs of the user, `modelbase` greatly simplifies the model-building process, by facilitating a systematic construction of kinetic models fully embedded in the Python programming language, and by providing a set of functionalities that help to conveniently access and analyse the results.

Despite the fact that mathematical models vary significantly in their complexity, from very simple and abstract models to extremely detailed ones, they share a set of universal properties. The process of building a kinetic model can be divided into a number of mandatory steps such as i) establishing the biological network structure (the stoichiometry), ii) defining the kinetic rate expressions, iii) formulation of the differential equations, iv) parametrisation, v) validation and, finally, vi) application [1]. `modelbase` supports researchers in every step of model development and application with its simple design aimed at being minimally restrictive. It has been written in Python, an open source, general-purpose, interpreted, interactive, object-oriented, and high-level programming language. Due to a long list of its general features, such as clear syntax, useful built-in objects, a wealth of general-purpose libraries, especially NumPy and SciPy, Python has become a widely used scientific tool [2]. Needless to say, the usage of Python over other, proprietary software such as MATLAB or Wolfram Mathematica, decreases the risk of limited reproducibility and transparency, two critical factors while conducting research. Unfortunately, several powerful models of central biochemical pathways [3, 4] have been published before this need became apparent. As a consequence, some of these models are extremely difficult to implement to even attempt to reproduce their results. Therefore, `modelbase` provides an environment for relatively easy implementation of models that were published without source code, in a general-purpose and reusable format. Moreover, `modelbase` supports the export of a structural (stoichiometric) model into Systems Biology Markup Language (SBML) for further structural analysis with the appropriate software.

In recent years, several other Python-based modelling tools have been developed, such as ScrumPy [5]; or PySCeS [6]. They allow performing various analyses of biochemical reaction networks, ranging from structural analyses (null-space analysis, elementary flux modes) to kinetic analyses and calculation of control coefficients. To the best of our knowledge they do not provide dedicated methods for model construction inside Python, and the standard usage relies on loading previously assembled model definition files.

The `modelbase` package presented here provides an alternative toolbox, complementing the functionalities of existing programs for computer modelling. Its power lies mainly in integrating the model construction process into the Python programming language. It is envisaged that `modelbase` will greatly facilitate the model construction and analysis process as an integral part of a fully developed programming environment.

### Motivation

In the course of our photosynthetic research, we identified several shortcomings that are not adequately met by available free and open source research software. When constructing a series of similar models, which share the same basic structure but differ in details, it is, in most modelling environments, necessary to copy the model definition file (or even pieces of code) and perform the desired modifications. This makes even simple tasks, such as changing a particular kinetic rate law, hideous and unnecessarily complicated, affecting the overall code readability. To facilitate a systematic and structured model definition, exploiting natural inheritance properties of Python objects, our intention was to fully integrate the model construction process into the Python programming language, allowing for an automated construction of model variants. The necessity for this fully Python-embedded approach became further evident for isotope label-specific models [7], where an automatic construction of isotope-specific reactions from a common rate law and an atom transition map is desired. Such models are, for example, required to explain complex phenomena, such as the asymmetric label distribution during photosynthesis, first observed by Gibbs and Kandler in the 1950s [8, 9].

### Implementation and architecture

`modelbase` is a console-based application written in Python. It supplies methods to construct various dynamic mathematical models, using a bottom-up approach, to simulate the dynamic equations, and analyse the results. We deliberately separated construction methods from simulation and analysis, in order to reflect the experimental approach. In particular, a model object constructed using the *Model* class can be understood as a representation of a model organism or any subsystem, on which experiments are performed. Instances of the *Simulator* class in turn correspond to particular experiments. The software components of `modelbase` are summarised in the UML diagram in Figure 1.

**Figure 1:**
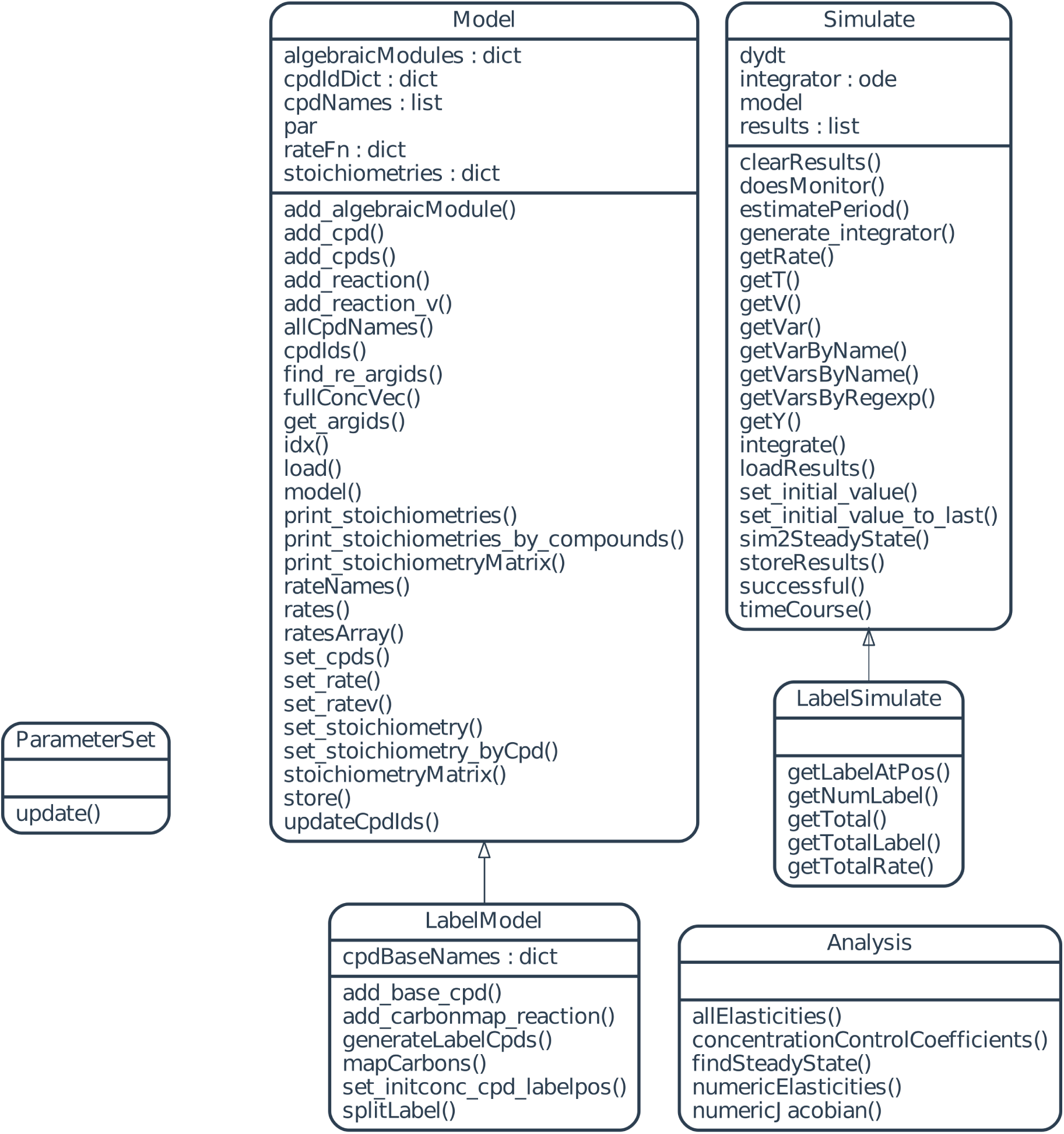
Diagrammatic representation of software components.

### Model construction

The user has the possibility to build two types of models, using one of the classes defined in the module *model*: *Model*, for differential-equation based systems, or *LabelModel*, for isotope-labelled models.

### Class *Model*

Every model object is defined by:

1. model parameters,
2. model variables,
3. rate equations,
4. stoichiometries.

Model parameters can be simply defined in a dictionary, `d`. To build a mathematical model the user needs first to import the `modelbase` package and instantiate a model object (called `m`).

**`import modelbase`**

`m = modelbase.Model(d)`

After instantiation, the keys of the parameter dictionary `d` become accessible as attributes of an object of the internal class `modelbase.parameters.ParameterSet`, which is stored as the model’s attribute `m.par`.

To add reacting entities of the described system (referred to as species in SBML), e.g., metabolites, we pass a list of compounds names to the `set_cpds` method:

`m.set_cpds(list_of_compounds)`

Each of the added compounds becomes a state variable of the system. The compounds are accessible by their name (string). The full list of all variables is stored in the attribute `m.cpdNames`.

If *S* denotes the vector of concentrations of the biochemical reactants (as defined with the method `set_cpds`), the temporal change of the concentrations is governed by

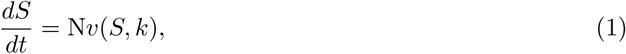

where N denotes the stoichiometric matrix and *v*(*S*, *k*) the vector of reaction rates as functions of the substrate concentrations *S* and parameters *k*. The system of ordinary differential equations is assembled automatically after providing all reaction rates and their stoichiometries to the method `m.add_reaction()`. The stoichiometric matrix of a model can be retrieved by the method `m.print_stoichiometries()` or `m.print_stoichiometries_by_compounds()`, for the transposed matrix. A detailed example of instantiating objects and solving a simple biochemical system with three reactions and two metabolites is provided in Box 1.

#### Box 1: Basic model use

We use `modelbase` to simulate a simple chain of reactions, in which the two state variables X and Y describe the concentrations of the intermediates. We assume a constant influx rate v_0_, a reversible conversion of X to Y, described with mass action kinetics with forward and backward rate constants k_1*p*_ and k_1*m*_, respectively, and an irreversible efflux of Y described by mass action kinetics with the rate constant k_2_.

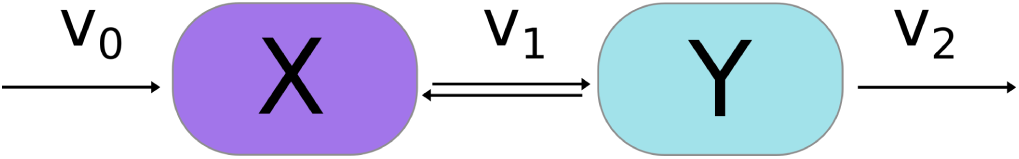

We import the `modelbase` package, define a list of metabolite species and a dictionary with parameters

~~~
import modelbase
cmpds = [’X’,’Y’]
p = {’v0’:1,’k1p’:0.5,’k1m’:1,’k2’:0.1}
~~~

We instantiate a model object of class *Model*

`m = modelbase.Model(p)`

and pass metabolites to the model (variables are always defined by names)

`m.set_cpds(cmpds)`

In the next step we define reaction rates functions. The rate functions always accept the model parameters as first argument, whilst the remaining arguments are metabolite concentrations.

~~~
v0 = lambda p: p.v0
def v1(p,x,y):
      return p.k1p*x - p.k1m*y
def v2(p,y):
      return p.k2*y
~~~

and then pass them to the model using `add_reaction()`

~~~
m.add_reaction(’v0’, v0, {’X’:1})
m.add_reaction(’v1’, v1, {’X’:-1,’Y’:1}, ’X’, ’Y’)
m.add_reaction(’v2’, v2, {’Y’:-1}, ’Y’).
~~~

To perform the computation we generate an instance of the class *Simulator*

`s = modelbase.Simulator(m)`

To integrate the system over a given period of time (`T=np.linspace(0,100,1000)`), with the initial concentrations set to 0 (`y0=np.zeros(2)`), we use the method `timeCourse()`

`s.timeCourse(T, y0).`

Convenient access to the results of simulation through various `get*()` methods enables easy graphical display.

~~~
plt.figure()
plt.plot(s.getT(),s.getY())
plt.legend(m.cpdNames)
~~~

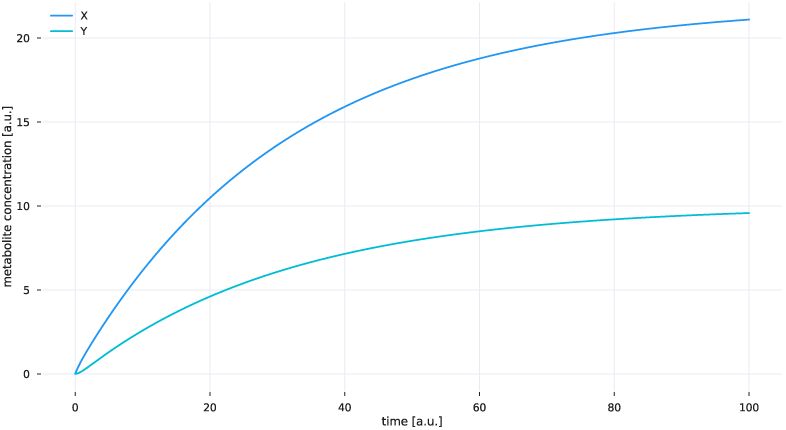

### Working with algebraic modules

A particularly useful function of the class *Model* has been developed to facilitate the incorporation of algebraic expressions, by which dependent variables can be computed from independent ones. Examples include conserved quantities, such as the sum of adenine phosphates, which is often considered to be constant, and rapid-equilibrium or quasi steady-state approximations, which are applicable for systems with time-scale separation and allow calculation of fast from slow variables. The function `add_algebraicModule()` accepts as arguments a function describing the rule how the dependent variables are calculated from independent ones, the name of the newly created module, and lists of names of the independent and dependent variables. After definition of an algebraic module, all dependent variables become directly accessible. The full list of independent and dependent variables can be accessed using the method `allCpdNames()`.

### Class *LabelModel* for isotope-labelled models

The `modelbase` package includes a class to construct isotope-labelled versions of developed models. In order to simulate the dynamic distribution of isotopes, `modelbase` defines dynamic variables representing all possible labelling patterns for all intermediates. In contrast to instances of the class *Model*, for instances of the class *LabelModel* for every compound the number of potentially labelled atoms (usually carbon) needs to be defined. This done with the method `add_base_cpd()`, which accepts the name and the number of labelled atoms of the compound. It automatically creates all 2^*N*^ isotope variants of the compound, where *N* denotes the number of labelled atoms. Finally, the method `add_carbonmap_reaction()` automatically generates all isotope-specific versions of a reaction. It accepts as arguments the reaction name, rate function, carbon map, list of substrates, list of products and additional variables to be passed.

`m = modelbase.LabelModel()`

With an instance of this class, for example the dynamic incorporation of radioactive carbon during photosynthesis can be easily defined and simulated, using the *Simulator* class described below. An example of how to use this class is provided in the Box 2.

### Integration methods and *Simulator* subpackage

The *Simulator* class of `modelbase` provides computational support for dynamic simulations. It is instantiated by

`s = modelbase.Simulator(m)`

and provides methods to numerically simulate the differential equation system and to analyse the results. Simple applications to run and plot a time course are given in boxes 1 and 2. By default, the dynamic equations are numerically integrated using a CVODE solver for stiff and non-stiff ordinary differential equation (ODE) systems. The default solver uses the Assimulo simulation package [10], with the most central solver group originating from the SUNDIALS (a SUite of Nonlinear and DIfferential/ALgebraic equation Solvers) package [11]. If Assimulo is not available, standard integration methods from the SciPy library [12] are used. When needed, almost every integrator option can be overridden by the user by simply accessing

`s.integrator`

Additionally, the *Simulator* class includes methods to integrate the system until a steady-state is reached (`sim2SteadyState()`), and to estimate the period of smooth limit cycle oscillations (`estimatePeriod()`). The solution arrays are accessed with the methods `getT()` and `getY()`. The advantage of using this method over using Assimulo’s `integrator.ysol` is that `getY()` also returns the result for all the derived variables (for which algebraic modules have been used). The powerful python plotting library matplotlib [13] provides numerous methods for graphical display of the results.

#### Box 2: Isotope-labelled model

A minimal example of an isotope-label specific model simulates equilibration of isotope distribution in a system consisting of the two reactions of triose-phosphate isomerase and fructose-bisphosphate aldolase:

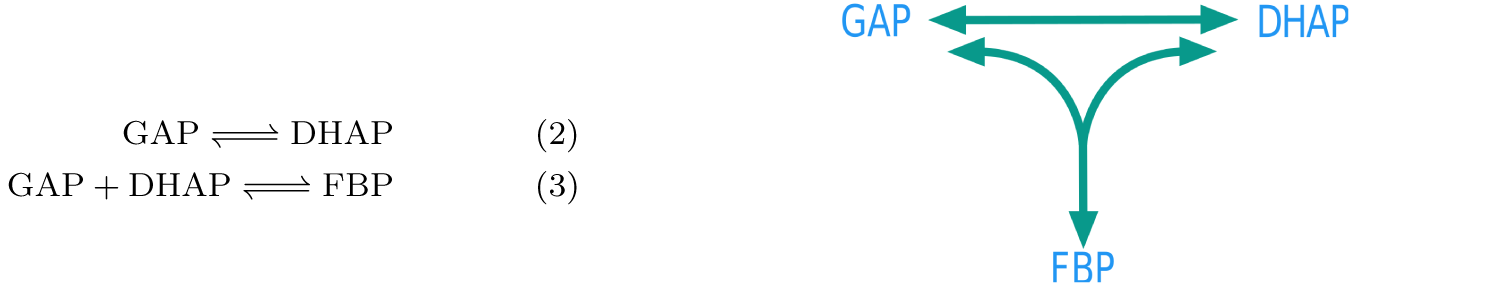

We define a dictionary of parameters and instantiate the model of class *LabelModel*

~~~
p={’kf_TPI’: 1,’Keq_TPI’: 21,’kf_Ald’: 2000,’Keq_Ald’: 7000}.
m = modelbase.LabelModel()
~~~

Compounds are added with an additional argument defining the numbers of carbons

~~~
m.add_base_cpd(’GAP’, 3)
m.add_base_cpd(’DHAP’, 3)
m.add_base_cpd(’FBP’, 6)
~~~

leading to an automatic generation of 80 = 2^6^ + 2^3^ + 2^3^ isotope-specific compounds. All reactions are assumed to obey mass-action rate laws. Standard rate laws are defined in the `modelbase.ratelaws` module.

~~~
import modelbase.ratelaws as rl
def v1f(p,y):
  return rl.massAction(p.kf_TPI,y)
~~~

All isotope-specific rates are generated by the `add_carbonmap_reaction()` method, based on a list defining in which positions the carbons appear in the products.

~~~
m.add_carbonmap_reaction(’TPIf’,v1f,[2,1,0],[’GAP’],[’DHAP’],’GAP’).
~~~

We set the initial conditions such that the total pools are in equilibrium, but carbon 1 of GAP is fully labelled

~~~
GAP0 = 2.5e-5
DHAP0 = GAP0 * m.par.Keq_TPI
y0d = {’GAP’: GAP0,
              ’DHAP’: DHAP0,
              ’FBP’: GAP0 * DHAP0 * m.par.Keq_Ald}
y0 = m.set_initconc_cpd_labelpos(y0d,labelpos={’GAP’:0})
~~~

and simulate equilibration of the labels for 20 arbitrary time units

~~~
s = modelbase.LabelSimulate(m)
T = np.linspace(0,20,1000)
s.timeCourse(T,y0)
~~~

and plot the result using the `getLabelAtPos()` method.

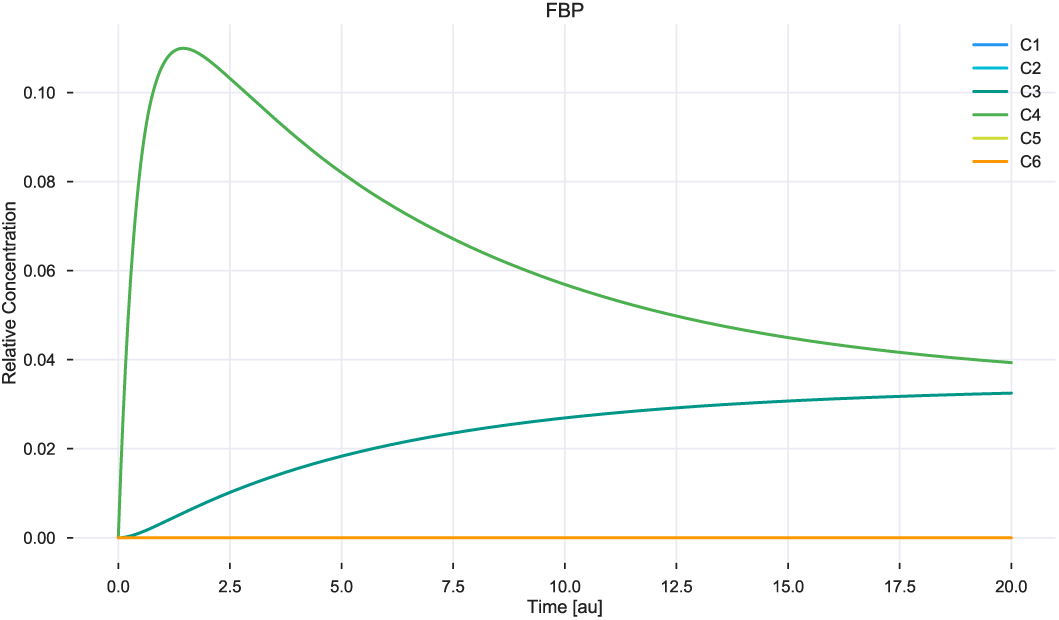

### Simple analysis with *Analysis* subpackage

The class *Analysis* of `modelbase` provides more advanced analysis methods complementing the simulations. Currently, it provides methods to numerically calculate elasticities and the Jacobian, find steady states by attempting to solve the algebraic equations, and to calculate concentration control coefficients. We expect the range of analysis methods to increase continuously in the future.

### Systems Biology Markup Language (SBML)

`modelbase` supports export of a structural (stoichiometric) version of a created model into an XML file in the computer-readable SBML format. Structural and stoichiometric analyses are currently not implemented in `modelbase`, therefore such export allows to take advantage of other SBML compatible modelling environments developed for such tasks (e.g. PySCeS or CobraPy [14]). The import of SBML models into `modelbase` is currently not supported, mainly because of the complementary purpose for which it was developed. The `modelbase` framework simplifies construction of kinetic models, allowing to perform this task with minimal modelling experience. Therefore, the main purpose of `modelbase` is the model design process itself, rather than importing a predefined construct to perform complex computations. However, a full SBML export and import functionality is currently under development to allow model sharing across different environments and platforms.

### Quality control

`modelbase` has been continuously developed and used within our lab since 2016. It has been successfully applied to study the complexity of photosynthesis and carbon assimilation in plants [7] and is being further maintained and developed.

## (2) Availability

### Operating system

`modelbase` is compatible with all platforms with working Python distribution.

### Programming language

`modelbase` is written in the Python programming language, a general-purpose interpreted, interactive, object-oriented, and high-level programming language. It is available for every major operating system, including GNU/Linux, Mac OSX and Windows and has been tested with Python 3.6.

### Additional system requirements

None

### Dependencies

Dependencies are provided in the `setup.py` file and include:

- numpy 1.14.3
- scipy == 1.1.0
- numdifftools == 0.9.20
- assimulo ==2.9
- pandas == 0.22.0
- python-libsbml == 5.17.0

Support for the differential equation solver sundials (CVODE) through the python package assimulo requires moreover

- Sundials-2.6.0 (for 64bits machines, install Sundials using ‐fPIC)
- Cython 0.18
- C compiler
- Fortran compiler

### List of contributors

In alphabetic order: Marvin van Aalst, Oliver Ebenhoh, Anna Matuszyhska, Nima Saadat.

### Software location

**Archive**

**Name:** Python Package Index (PyPI)
**Persistent identifier**: https://pypi.org/project/modelbase/
**Licence:** GPL4
**Publisher:** Oliver Ebenhöh
**Version published:** 0.2.3
**Date published:** 25/06/18

**Code repository**

**Name**: GitHub
**Persistent identifier:** https://github.com/QTB-HHU/modelbase
**Licence:** GNU General Public License v3.0
**Date published:** 25/06/18

### Language

`modelbase` was entirely developed in English.

## (3) Reuse potential

The strength of our package lies in its flexibility to be applied to simulate and analyse various distinct biological systems. It can be used for the development of new models, as well as reconstruction tool. We demonstrate its power by reconstructing three mathematical models that have been previously published without providing the source code (Table 1). This includes a model of the photosynthetic electron transport chain, originating from our lab, that initially has been developed in MATLAB [15].

**Table 1:**
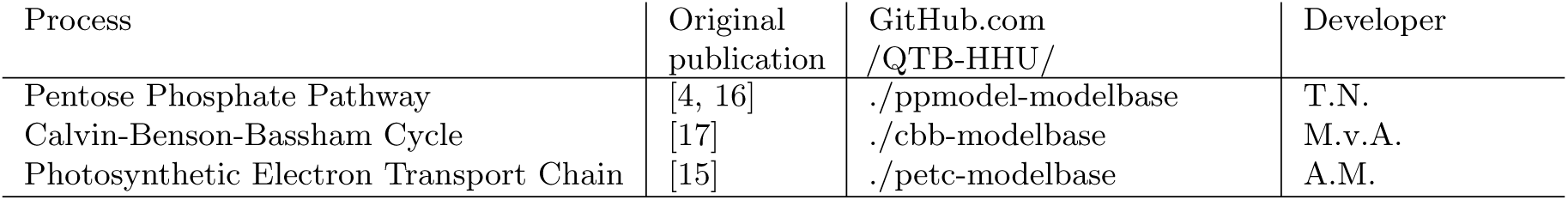
Mathematical models originally published without the source-code, reconstructed in our lab using the `modelbase` package. The source code and examples are available from the GitHub repository of our lab https://github.com/QTB-HHU/

### Modelling photoprotective mechanisms

Part of our research focuses on understanding the dynamics of various photoprotective mechanisms present in photosynthetic organisms [15, 18, 19]. The foundation of our further work constitutes the model of the photosynthetic electron transport chain in green algae *Chlamydomonas reinhardtii* published in 2014 [15]. We have reimplemented the original work in Python and reproduced the results published in the main text (Figure 2), providing a photosynthetic electron transport chain core, compatible with other `modelbase`-adapted modules, to further our studies on the dynamics of light reactions of photosynthesis.

**Figure 2:**
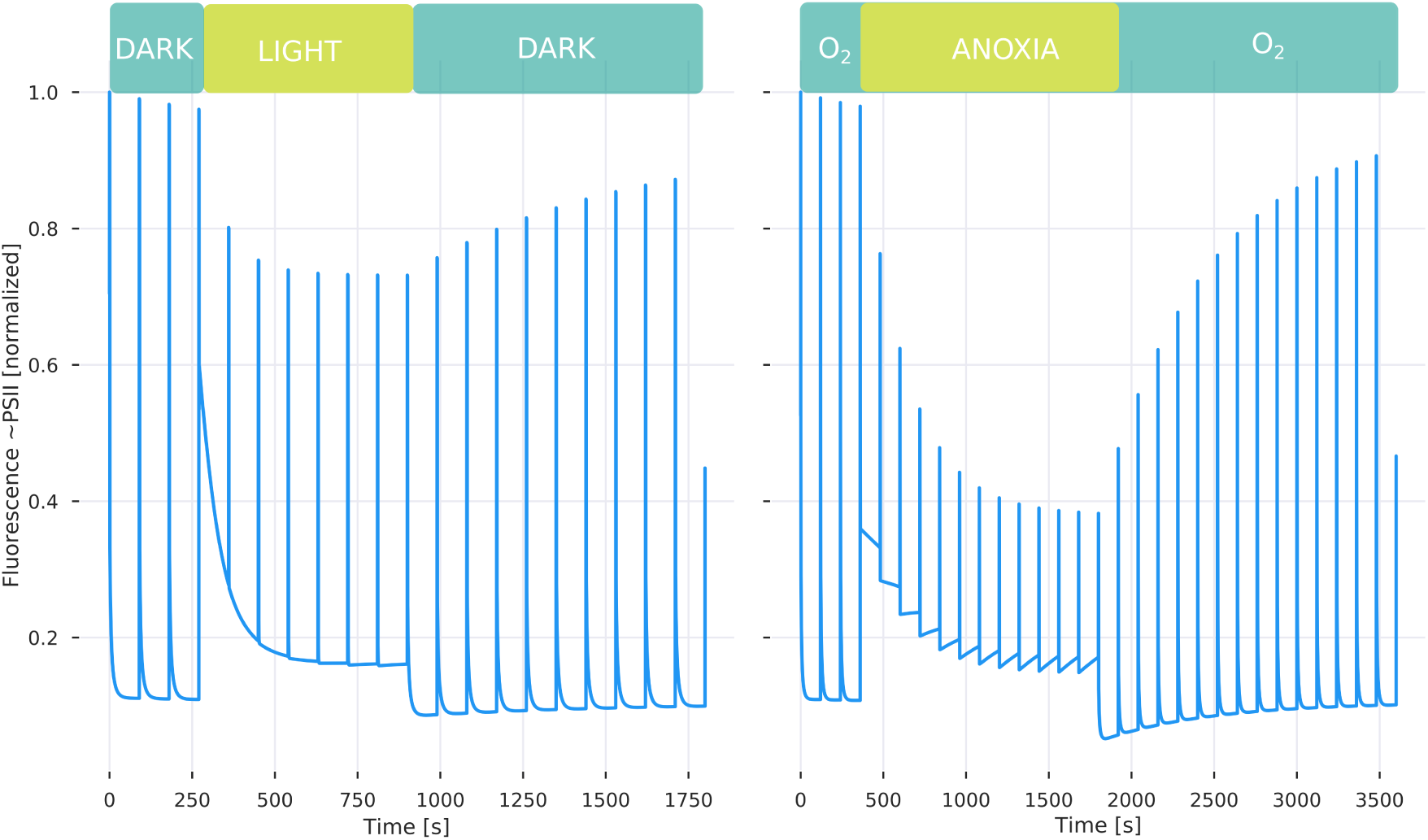
Reproduction of the Figures from the [15]. Simulated fluorescence trace obtained through Pulse Amplitude Modulation (PAM) under light induced (left) and anoxia induced (right) conditions. The dynamics of the fluorescence decrease corresponds to the activation of a specific photoprotective mechanism called state transitions, while the increase in the signal after the inducer (light or anoxia) is switched off relates to the relaxation of the mechanism.

### Dynamics of the carbon assimilation

Using `modelbase`, we have re-implemented a model of the Calvin-Benson-Bassham (CBB) Cycle by Poolman et al. [17]. The model is a variant of the Pettersson and Pettersson [3] model, where the strict rapid equilibrium assumption is relaxed and fast reactions are modelled by simple mass action kinetics. Its main purpose is to study short to medium time scale responses to changes in extra-stromal phosphate concentration and incident light. The concentrations of NADPH, NAD^+^, CO_2_ and H^+^ are considered constant, leaving the 13 carbohydrate cycle intermediates, ATP, ADP and inorganic phosphate as dynamic variables. The model further incorporates a simplified starch production using glucose 6-phosphate and glucose-1-phosphate and a simple ATP recovery reaction. We used the `modelbase` implementation of the Poolman model to simulate the steady state concentrations of the metabolites depending on the extra-stromal phosphate concentration (Figure 3), reproducing original work by Pettersson and Pettersson [3]. We have observed that the system is not stable any more for [*P*_*ext*_] > 1.5, a feature not discussed in the Poolman paper [17].

**Figure 3:**
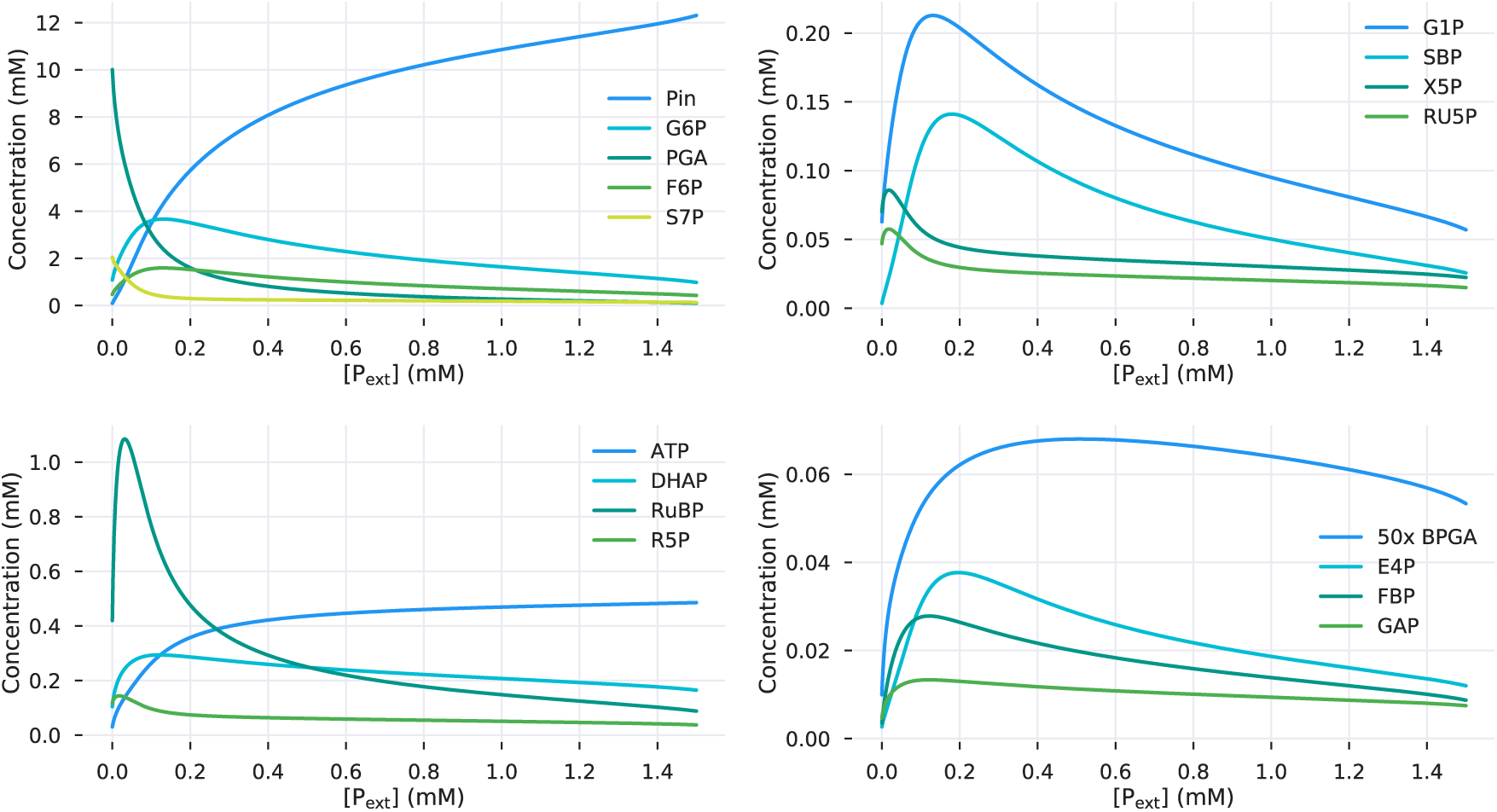
Metabolite steady state concentrations dependent on the extra-stromal phosphate concentration simulated with the Poolman implementation of the Pettersson and Pettersson model of the CBB cycle [17].

The compatible mathematical representation of the two photosynthetic subsystems, the ATP-producing light reactions and the ATP-consuming CBB cycle, is a prerequisite to merge those two models. Technically, in the `modelbase` framework, this is a straight forward process. Scientifically, it turned out to be not a trivial task (unpublished work).

### Pentose phosphate pathway

We envisage that especially our *LabelModel* extension will find a wide application in metabolic network analysis. Radioactive and stable isotope labelling experiments constitute a powerful methodology for estimating metabolic fluxes and have a long history of application in biological research [20]. Here, we showcase the potential of `modelbase` for the isotope-labelled experiments by reimplementing the model of the F-type non-oxidative Pentose phosphate pathway (PPP) in erythrocytes originally proposed by McIntyre *et al*. [4]. This was later adapted by Berthon *et al*. for label experiments and *in silico* replication of ^13^Carbon nuclear magnetic resonance (NMR) studies [16]. We have reproduced the results obtained by the authors, including the time course of diverse Glucose-6-phosphate isotopomers (Figure 4).

**Figure 4:**
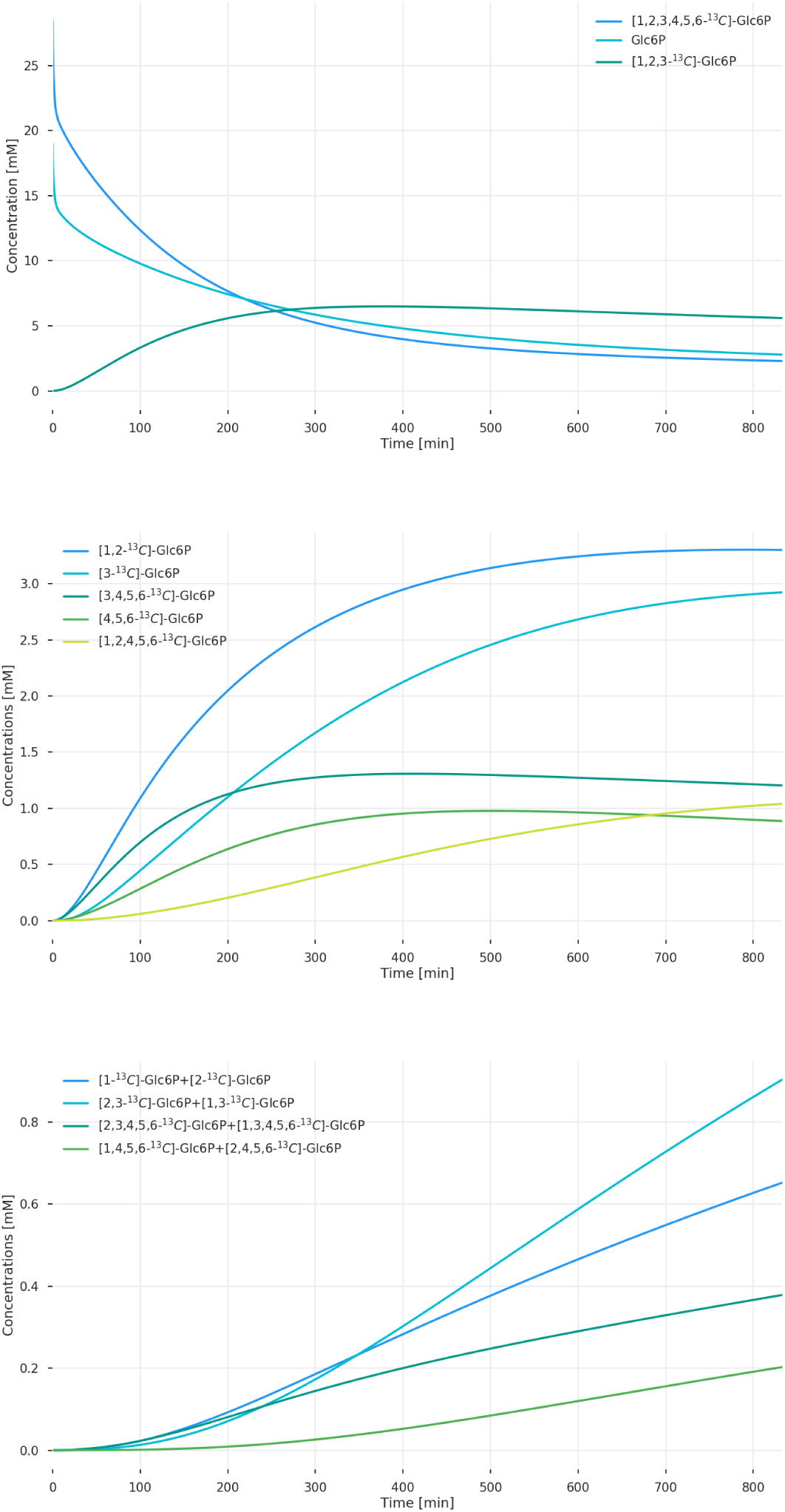
Formation of diverse Glc6P isotopomers in a haemolysate, obtained by solving the adapted model by Berthon *et al*. [16] reimplemented using `modelbase`.

Among many other applications, `modelbase` provides tools to reproduce the ‘photosynthetic Gibbs effect’. Gibbs and Kandler described it in 1956 and 1957 [8, 9], when they observed the atypical and asymmetrical incorporation of radioactive ^14^CO_2_ in hexoses. An example of label incorporation by the CBB cycle intermediates is presented schematically in Figure 5.

**Figure 5:**
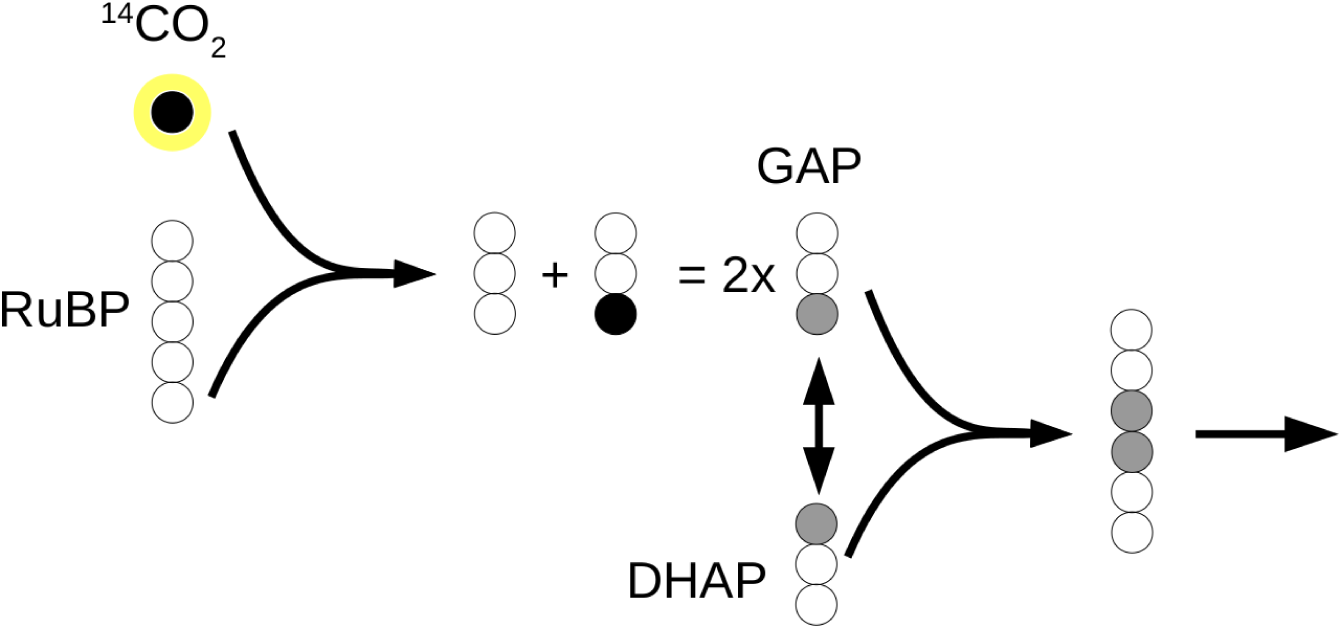
Schematic representation of label incorporation by the CBB cycle intermediates.

Finally, our package provides a solid foundation for additional extensions to the framework architecture, its classes and modelling routines. To encourage its use and to facilitate the first steps to apply the `modelbase` package, we have prepared an interactive tutorial using a Jupyter Notebook [21], which showcases basic implementation of `modelbase` and each of its classes in easy to follow and thoroughly explained examples (see https://github.com/QTB-HHU/modelbase/tutorial.ipybn).

## Acknowledgements

We would like to thank the students working on their bachelor and master projects in our lab, who applied and tested this software while investigating their scientific problems.

## Funding statement

This work was financially supported by the Deutsche Forschungsgemeinschaft “Cluster of Excellence on Plant Sciences” CEPLAS (EXC 1028).

## Competing interests

The authors declare that they have no competing interests.

## Paper Author Roles

O.E. initiated the project, developed the code and provided teaching examples; M.v.A. and A.M developed further the code; M.v.A. reimplemented the Calvin-Benson-Bassham Cycle model as an example of `modelbase` utility, T.N. reimplemented the Pentose-Phosphate-Pathway model and A.M. reimplemented the photosynthetic electron transport chain model; N.P.S. provided export support for SBML models; A.M. prepared the Jupyter Notebook with the tutorial and wrote the first draft of the manuscript. All authors have read the manuscript and contributed to its final version.

## Abbreviations

CBB: Calvin-Benson-Bassham
NMR: Nuclear Magnetic Resonance
ODE: Ordinary Differential Equations
PAM: Pulse Amplitude Modulation
PPP: Pentose-Phosphate
SBML: Systems Biology Markup Language
UML: Unified Modeling Language

